# A large-scale assessment of lake bacterial communities reveals pervasive impacts of human activities

**DOI:** 10.1101/821991

**Authors:** S.A. Kraemer, N. Barbosa da Costa, B.J. Shapiro, Y. Huot, D. Walsh

## Abstract

Lakes play a pivotal role in ecological and biogeochemical processes and have been described as ‘sentinels’ of environmental change. Assessing ‘lake health’ across large geographic scales is critical to predict the stability of their ecosystem services and their vulnerability to anthropogenic disturbances. The LakePulse research network is tasked with the assessment of lake health across gradients of land use on a continental scale. Bacterial communities are an integral and rapidly responding component of lake ecosystems, yet large-scale responses to anthropogenic activity remain elusive. Here, we assess the ecological impact of land use on bacterial communities from 220 lakes covering more than 660 000 km^2^ across Eastern Canada. Alteration of communities was found on every level examined including richness, community composition, community network structure and indicator taxa of high or low lake water quality. Specifically, increasing anthropogenic impact within the watershed lowered richness mediated by changes in salinity. Likewise, community composition was significantly correlated with agriculture and urban development within a watershed. Interaction networks showed decreasing complexity and fewer keystone taxa in impacted lakes. Together, these findings point to vast bacterial community changes of largely unknown consequences induced by human activity within lake watersheds.

**Significance Statement:** Lakes play central roles in Earth’s ecosystems and are sentinels of climate change and other watershed alterations. Assessing lake health across large geographic scales is therefore critical to predict ecosystem stability and lake vulnerability to anthropogenic disturbances. In this context, the LakePulse research network is tasked with a large-scale assessment of lake health across Canada. Bacterial communities are an integral and rapidly responding component of lake ecosystems, yet their large-scale responses to anthropogenic activity remain unknown. Here, we assessed the anthropogenic impact on bacterial communities of over 200 lakes located across large environmental gradients. We found communities to be impacted on every level investigated, indicating that human activities within watersheds cause vast bacterial community changes of largely unknown consequences.

## Introduction

Lake ecosystems have garnered a large amount of attention from researchers and policy makers in recent years as ‘sentinels’ of climate change and human impacts (Adrian *et al.* 2009) owing to their central importance in biogeochemical cycles (Tranvik *et al.* 2009). This role arises from their pivotal position within watersheds from which they receive nutrients and environmental contaminants. Consequently, many anthropogenic activities in the watershed threaten the ecosystem and freshwater services provided by lakes. For example, intense agricultural activity, and specifically the use of fertilizers within watersheds, has been connected to lake eutrophication (Arbuckle & Downing 2001; Taranu & Gregory-Eaves 2008), in conjunction with the bloom of certain species of cyanobacteria (Heisler *et al.* 2008), decreases in oxygen conditions (Scavia *et al.* 2014), and increased methane emissions (Bastviken *et al.* 2004), while watershed urban development has been connected to overall water degradation caused by road salt runoff (Novotny *et al.* 2008; Dugan *et al.* 2017) and phosphorus export (Hobbie *et al.* 2017). However, many important human impacts with significant consequences for ecosystem function are likely undetected and therefore cannot be addressed by policy or monitoring programs.

Bacterial communities mediate essential biogeochemical processes within lakes but have historically been much less explored than other members of lake food webs such as fish and zooplankton. Recent advances in microbial molecular ecology have demonstrated high taxonomical and functional diversity of lake bacteria and links between shifts in community composition and activity and ecosystem-level responses (Shade *et al.* 2007; Kara *et al.* 2013). Given the high bacterial diversity and ability to respond rapidly to changing environments, these communities may be powerful indicators of environmental stressors (e.g. excess nutrient loading, acidification, salinification, metal contamination). However, little is known about the variability of freshwater bacterial communities across large spatial gradients of land use (but see Marmen et al. 2019).

Canada is responsible for the stewardship of more than 1,000,000 lakes, which make up 20% of the world’s freshwater stocks and provide the main source of drinking water for many major Canadian cities (Environment Canada 2011; Huot *et al.* 2019). Lake ‘health’ has previously been described as the departure of the ecological state of a lake from the pristine state; the further removed from the pristine state the less healthy a lake is (Huot et al. 2019). While large-scale assessments of lake health have been conducted elsewhere (Hering *et al*. 2013, US EPA 2009), no such survey exists for Canadian freshwaters. In this context, the LakePulse network has embarked upon a continental-scale assessment of lake health across Canada in relation to anthropogenic activity as determined from satellite imaging and lake environmental variables collected in situ. In this study, we connect the bacterial community structure to watershed agriculture, forestry, pasture and urban development in more than 200 lakes covering over 660 000 km^2^ in Eastern Canada (Fig. 1A). We aim to untangle the underlying proximate environmental variables that cause the community shifts observed, as well as describe how anthropogenic impacts change bacterial interaction networks. Lastly, we use the scope of our dataset to investigate the relative strengths of different processes of community assembly such as drift and selection. We found that communities were markedly altered with respect to both their richness and community composition by watershed anthropogenic activity, specifically agriculture and urban development, which increased lake salinity. Moreover, highly impacted watersheds harboured more fragmented bacterial communities with fewer keystone taxa, which could affect ecosystem function and resilience. Although the study will be extended across Canada over the next few years, we were compelled by the observation of deep alterations in lake bacterial communities by human activities in a relatively less impacted region of Canada to report rapidly on these findings.

**Figure 1:**
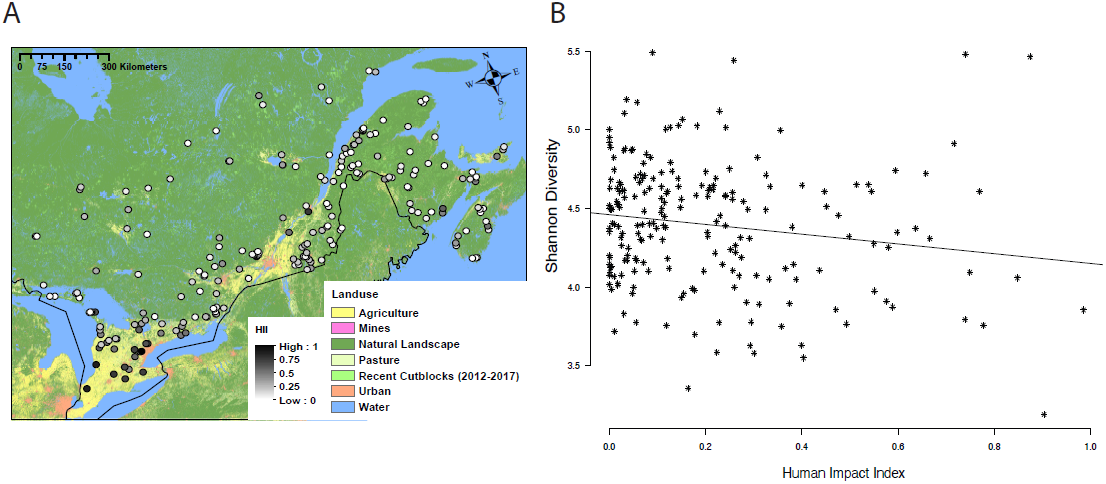
A) Map of Eastern Canada showing 210 sampled lakes coloured by their Human Impact Index. B) Scatterplot showing the relationship between Shannon diversity and the Human Impact Index variable for 210 lakes.

## Methods

### Lake selection

Lakes were selected across Eastern Canada as described in detail in Huot *et al.* (2019). Briefly, lakes were picked in a random sampling design with ecozone, lake size and human impact index as stratifying factors. Canadian ecozones (represented by AH: Atlantic Highlands, AM: Atlantic Maritimes, BS: Boreal Shield and MP: Mixedwood Plains in south-eastern Canada) represent regions with distinct geological, climatic and ecological features (Ecological Stratification Working Group 1996). For each lake within 1 km of a road the watershed was delineated as described in Huot *et al.* (2019). Within the watershed, each pixel was assigned a human impact value between 0 and 1 depending on their category (Urban development/Road: 1; Mines/Oil: 1, Agriculture: 1, Pasture: 0.5, Forestry (recent clear-cuts): 0.5, Natural landscapes: 0) and the average Human Impact Index (HII) for the lake calculated across the watershed. Across the ecozones of Eastern Canada, the Mines/Oil category was found to be negligible in magnitude (only five lakes total with > 1% mining within their watershed and only one lake with > 5% mining) and was thus not explicitly considered in the analyses of specific land use classes herein.

### Surface water sampling

The surface water of 220 lakes (Fig. 1A) was sampled during the time of maximum summer lake stratification between July and September 2017 at the deepest point of each lake (as measured on site using a sonar). The depth of the euphotic zone was determined as twice the Secchi disk depth and an integrated epilimnion sampler (sampling tube) was used to sample to that depth, or to a depth of 2 m, whichever was shallower. Water samples were immediately transferred and stored in the cold and dark, subsequently filtered through a Durapore 0.22 μm membrane filter (Sigma-Aldrich, St. Louis, USA) on site and immediately frozen at −80°C until analysis. In addition, for each lake we used the same water sample to determine dissolved organic and inorganic carbon (DOC and DIC (mg/L), supplemental material), total nitrogen (TN (mg/L), (Patton & Kryskalla 2003)), concentrations of the ions magnesium, potassium, sodium (U.S. Environmental Protection Agency 1994), chloride and sulfate (U.S. Environmental Protection Agency 1997) (all in mg/L). The average concentration of dissolved oxygen (DO, mg/mL) over the same sampling depth were measured by averaging the values from a multiparameter probe profile (RBR Maestro^3^ profile, RBR Ltd., Ottawa, Canada; with a fast oxygen probe Rinko III, JFE Advantech Co., Nishinomiya, Japan) over that depth.

### Processing of environmental and spatial data

Distances between the lakes (km) were used to calculate Moran’s eigenvector maps (MEMs, *adespatial* R package; Dray *et al.* 2016). We visually inspected the MEM eigenvalue distribution and chose the first six MEMs to describe spatial structure within our system. We extracted the six MEMs’ coordinates and combined them with data on the lake area, lake depth, DOC, DIC, TN, DO, magnesium, potassium, sodium, chloride and sulfate. Lakes with missing data for any of these variables were removed and a PCA (after scaling of the environmental variables) performed for the remaining 167 lakes (*rda* function, *vegan* R package, Oksanen *et al.* 2016). The first seven principle component axes (PCs) were chosen for further analysis as they represented more variation than the mean. Factor loadings for these PCs are shown in Supplemental Table 1 and we considered factor loadings >=|0.43| to be significant (Hair *et al.* 1998). Ecozone, HII and size class membership, as well as environmental data for each lake can be found in Supplemental Table 2.

### DNA extraction and sequencing

DNA was extracted from filters with PowerWater kits (Mobio Technologies Inc., Vancouver, Canada) and the V4 region of the 16S rRNA gene PCR-amplified using the standard primers U515_F and E786_R before sequencing on an Illumina MiSeq machine. DNA extraction and sequencing succeeded for 212 of the 220 samples. Details of PCR and sequencing reactions can be found in the supplemental material.

### Sequence data processing

Read files were demultiplexed using idemp (Wu 2014) and Illumina adapters removed with cutadapt (Martin 2011). Reads were processed using the *DADA2* package in R (Callahan *et al.* 2016): Reads were trimmed, merged, chimeras removed and taxonomy of Amplicon Sequence Variants (ASVs) assigned to the genus level using the Fresh Train taxonomic database (Rohwer *et al.* 2018) and the SILVA database (version v128align) (Quast *et al.* 2013) to obtain an ASV table. Further analysis of the ASV table was done using the *phyloseq* package in R (McMurdie & Holmes 2013). ASVs from chloroplasts were removed before rarifying the data to a common read depth of 15000 reads after removing two lakes with less than 15000 reads. A phylogeny of the remaining 11510 ASVs was created using FastTree (Price *et al.* 2010). Bacterial richness of each lake was calculated using the Shannon index as implemented within the *phyloseq* package.

### Linear modeling of richness and community composition analyses

We modeled the impact of the Human Impact Index (HII) variable and of percent land use types in the watershed on bacterial richness using both generalized least squares (GLS models; *nlme* R package Pinheiro *et al.* 2019)) and linear mixed models (LMM; *lme4* package (Bates *et al.* 2015)). Linear mixed models additionally contained ecozone as a random effect, but either fitted worse than the corresponding GLS model or failed to converge.

We performed structural equation modeling (*lavaan* R package (Rosseel 2012)). We considered the impact of HII onto the first seven environmental PCs, as well as the impact of the PCs and HII directly onto ranked richness. Additionally, we modeled the impact of land use classes on the environmental PCs directly (supplemental material). Models were only considered to fit better than the null model if the chi-square *p*-value was > 0.05 and the comparative fit index > 0.95.

We utilized a PERMANOVA test to investigate the impact of the environmental PCs onto community composition (*adonis* function; *vegan* R package (Oksanen *et al.* 2016)). In addition, we performed forward selection db-RDA to determine how MEMs and watershed land use impact community composition (*capscale* function; *vegan* R package (Oksanen *et al.* 2016)).

### Networks of lake bacterial communities

Lakes were assigned to either a low (HII 0-0.1, 91 lakes), medium (HII>0.1&≤0.5, 96 lakes), or highly impacted (HII>0.5, 23 lakes) class. Within each network dataset, we excluded ASVs that were not present in at least 10 % of the samples, resulting in 826 ASVs in low-HII lakes, 775 ASVs in medium-HII, and 764 ASVs in high-HII lakes. We followed a combinatorial approach to determine significant connections between ASVs using both the Maximum Information Criterion (Albanese *et al.* 2013) and Sparse Inverse Covariance Estimation (Kurtz *et al.* 2015) as detailed in the Supplemental material.

We determined Order influence within high, moderate and low-impact networks by dividing the total number of edges of each Order by the number of nodes belonging to the Order (excluding Orders only represented by a single ASV or absent in any of the networks) (Banerjee *et al.* 2019). Moreover, we dropped all ASVs that could not be identified to the Order level.

### Indicator species of lake health

We investigated potential indicator species of high and low impact lakes based on abundance changes of taxa in pairs of lakes that were geographically similar but varied with respect to their HII. The log-ratio of ASV abundance change between paired lake communities was compared to control lake pairs. Additionally, we screened all ASVs for rank changes of at least 33% between paired impacted and pristine lakes as described in the supplemental material.

### Community assembly processes of lake bacterial communities

We defined generalist and specialist taxa within our dataset as follows: taxa with an occupancy ≥ 158 lakes (75% of the data set) were considered generalists, while ASVs in ≤ ten lakes (5%) and with a relative abundance of higher than 2% were considered specialists (Barberán *et al.* 2012). We investigated distance-decay curves between community dissimilarity (determined as the Bray-Curtis dissimilarity as calculated using the *vegdist* function in vegan (Oksanen *et al.* 2016), and the distance between the lakes (in km), using Mantel tests with 9999 permutations.

We followed the null model method as proposed by Stegen et al. (2013) to determine quantitative estimates for the strength of selection, dispersal limitation and ecological drift in the lake communities on a subsetted dataset in which rare taxa (abundance <500) were removed (679 ASVs remaining) as detailed in the supplemental material.

## Results

### Land use impact on bacterial richness, community composition and interactions

We firstly investigated the interaction between the human impact index (HII) and bacterial richness (here measured as the Shannon-Weaver index) of lake communities. Overall, we detected a significantly negative impact of HII on richness (slope: −0.31, *p*<0.05, Fig. 1B). When modeling richness as a function of different land use classes, we found the percentage of forestry within a watershed to have a significantly positive impact (slope: 1.26, *p*<0.01), while the percentage of urban development had a marginally significant negative impact (slope: −0.67, *p*=0.057).

To further investigate the relationship between HII and richness we performed structural equation modeling (SEM) to investigate the relationship between HII and the first seven environmental PCs and in turn the impact of the PCs onto ranked richness (Fig. 2). The SEM (χ^2^=5.684, DF=21, *p*=1.00, CFI=1.00) showed a significant positive impact of PC2 (*p*<0.001) and PC6 (*p*=0.006) on richness. PC2 is negatively loaded with lake depth, while PC6 carries the spatial eigenvector MEM2. PC6 was in turn significantly positively impacted by the HII variable (*p*=0.031). In addition, we detected a marginally significant negative impact of PC1 (*p*=0.056) on richness. While PC1 did not carry significant loadings, it is strongly negatively loaded with the ions Mg^+2^, K^+^, Ca^+2^ and Cl^−^. PC1 was strongly positively impacted by the HII variable (*p*<0.001). Lastly, the HII variable also directly impacted ranked richness (*p*=0.008).

**Figure 2:**
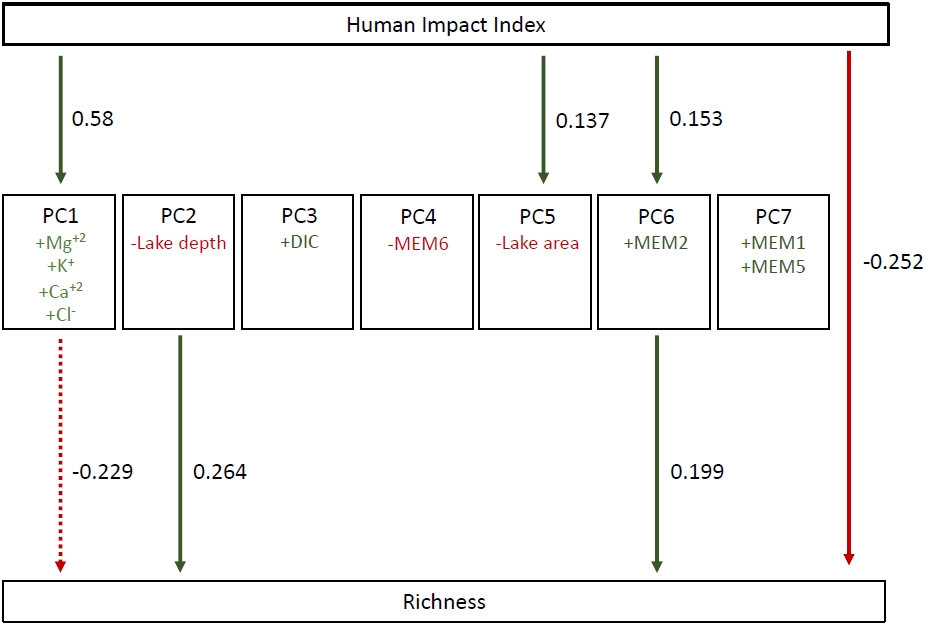
Graphical representation of the structural equation model. All paths were tested, but only those showing significant (p<0.05, solid lines) or marginally significant (p<0.1, dashed lines) correlations are shown. Positive interactions and loadings are shown in green (lighter green for high but non-significant loadings), negative in red.

Thus, HII was associated with increased lake ion concentrations, which in turn led to a decrease in bacterial richness (Fig. 2). Moreover, richness was impacted or correlated with lake depth. The link between HII, PC6 (loaded with a spatial eigenvector) and richness may indicate correlation with an unmeasured environmental variable. Lastly, HII also negatively impacted richness independent of the environmental PCs considered here, suggesting unknown environmental variables.

To understand the cause of the impact of the HII variable, we examined the underlying land use classes and investigated their relationship with ranked richness and the seven environmental PCs in an additional SEM (Supplemental material, Supp. Fig. 1). We found PC1 to be significantly positively associated with agriculture, urban development and pasture within the watershed.

We investigated factors structuring community composition via PERMANOVA with the first seven environmental PCs as explanatory variables. All PCs except PC7 were found to significantly impact community composition (all *p*<0.05). PC1, carrying ion concentration loadings, showed the highest R^2^ value in the model (0.062), but all factors included in the model only explained a small amount of the total variation (~18%).

We utilized db-RDA to select watershed-scale variables impacting community composition. A full model containing the spatial eigenvectors, lake morphometric parameters (lake depth and area) and land use data was found to be significant (*p*=0.001) so we proceeded with forward selection. Overall, a model containing the land use classes: natural landscapes, forestry, agriculture, pasture and urban development in the watershed, as well as the first four MEMs, and lake depth was selected as the best model (Fig 3). However, the model only explained ~15% of the variation observed.

**Figure 3:**
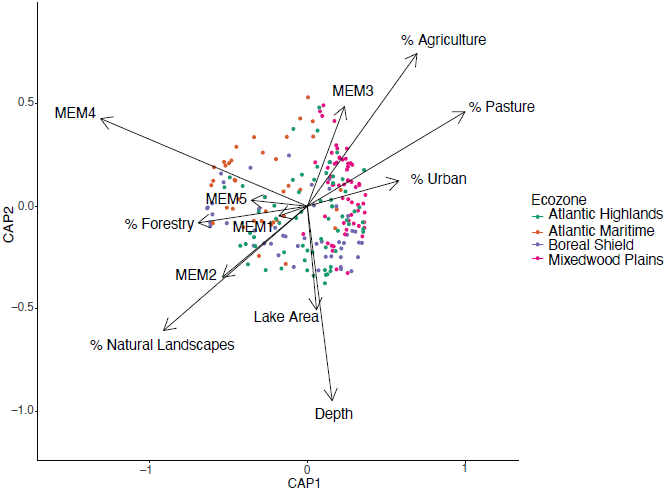
Plot of the db-rda coordinates of lake communities. Arrows show the significant environmental variables after forward selection.

To determine how anthropogenic impact may alter the structure of bacterial interactions, we constructed networks of high, moderate, and low HII lake communities. Low HII lake communities showed significant co-occurrence patterns for 179 nodes (taxa) with a total of 193 edges (co-occurrence connections). Moderate impact communities had significant co-occurrence networks for 145 nodes and 163 edges, while high impact communities had 220 nodes and 174 edges. Networks were more fragmented under high human impact (low: 24 components, moderate: 13 components, high: 59 components) and the clustering coefficient as well as the centralization value decreased with increasing HII (Fig. 4). Both low and moderate impact lake communities were characterized by ASVs having similar average numbers of neighbors (low: 2.16, medium: 2.25) and keystone taxa (with over five connections each: low: 13, moderate: 11), whereas ASVs in highly impacted lake communities had on average fewer neighbours (1.58). Lastly, highly impacted lakes only had two keystone taxa (taxa with >5 edges, Fig. 4).

**Figure 4:**
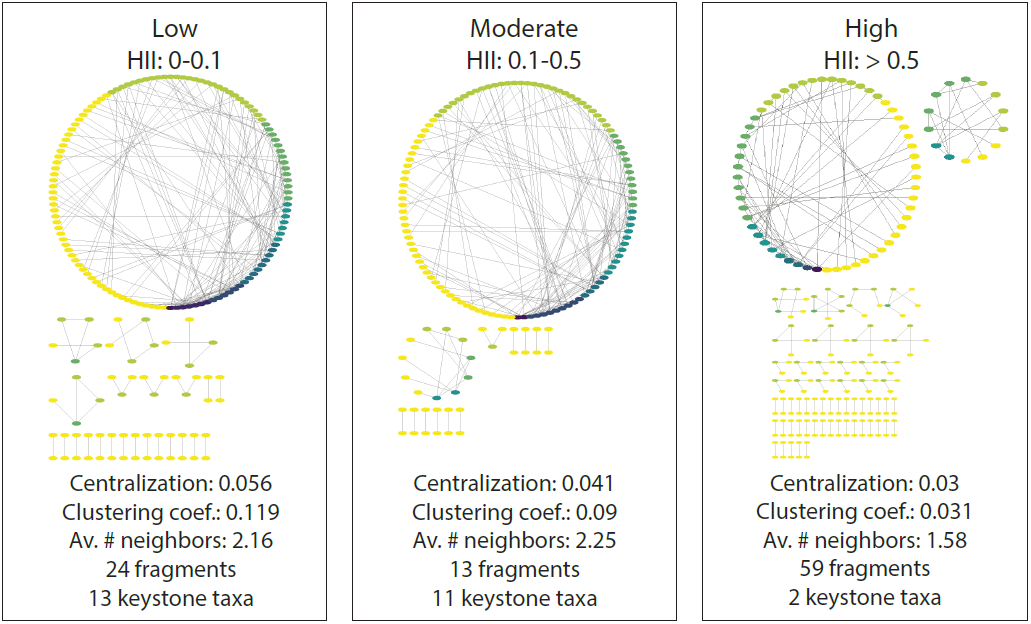
Co-occurrence networks and statistics of high, moderate and low impact lake communities. Nodes are colored by the number of their edges.

We determined whether specific bacterial Orders varied in their influence in high, moderate or low HII lakes by dividing their number of total edges by the number of nodes of a given Order. We identified 17 Orders which were represented in all three sets of networks (Supp. Table 3). Four of these Orders changed consistently between low, moderate and high lake networks. Burkholderiales ASVs, Frankiales ASVs and Rhizobiales ASVs had decreasing influence from low to moderate to high impacted networks, while verrucomicrobial OPB35’ASVs’ influence increased. Furthermore, Acidimicrobiales and Planctomycetales ASVs were of relatively low influence in high impact networks, but of higher influence in moderate and low impacted networks.

We investigated potential indicator species, i.e. species that can be used as ecological indicators for specific environmental conditions due to their niche preferences, for high or low impact lakes using log-ratio and rank abundance changes of ASVs. Overall, no strong indicator patterns emerged, demonstrating the high variability of freshwater communities under both pristine and impacted conditions. Log-ratio changes in treatment (pristine-impacted), but not control (pristine-pristine or impacted-impacted), lake pairs were detected for six ASVs in AH, two in AM, nine in BS and ten in MP. In total, 26 ASVs showed significant log-ratio changes, but these ASVs rarely overlapped between ecozones. Taking into account ASV taxonomy, we found ASVs belonging to the Burkholderiales *betI-A* clade and of the cyanobacterial *Family I* clade to be associated with low impact lakes across three and two ecozones, respectively (Supp. Table 4).

95 ASVs were found to show significant rank abundance changes between high and low impact lakes, but not between control lake pairs (17 ASVs in AH, 19 in AM, 46 in BS, 25 in MP, Supp. Table. 5). Specific taxa were associated with changing lake conditions across ecozones. For example, the Burkholderiales *betI-A* clade (represented by six ASVs) was associated with low impact lakes in all four ecozones, except for one ASV, which was a high impact indicator in AH.

### Evolutionary and ecological processes shaping lake bacterial communities

We did not detect any low-occupancy, high-abundance specialists within our dataset. In contrast, we identified 25 generalists, which were present in 75% or more of lakes. The most common generalist lineage was the actinobacterial *acI* lineage (9 ASVs), followed by the betaproteobacterial *betI* lineage (5 ASVs). Other lineages represented included *acIV* (2 ASVs), *betIV* (2 ASVs) and bacteroidal *bacI* (2 ASVs) (Supp. Table 6).

Overall, we found bacterial communities to be less similar the further apart in space they were sampled (Mantel test, *p*<0.001). Specifically, across all communities investigated, lakes sampled up to approximately 350 km apart were significantly more similar than expected by chance (Mantel correlogram analysis, *p*=0.004, Supp. Fig. 2A). To determine the causes for this structure, we performed a quantitative analysis of the community assembly processes shaping lake communities to estimate the relative strength of selection and drift processes in our dataset. Firstly, we determined whether we could detect a phylogenetic signal within our data (i.e. whether closely related species had similar niches) by testing for correlation of phylogenetic distance, based on part of the 16S rRNA gene, with ecological distance (differences in abundance patterns expressed as Bray-Curtis dissimilarity). Overall, we found a significant decay of phylogenetic distance with ecological distance (Mantel test: *p*<0.05, Supp. Fig. 2B). Specifically, and as expected, we detected a phylogenetic signal over relatively short phylogenetic distances (cophenetic distance>0.365), indicating that the beta nearest taxon index (βNDI), which indicates the impact of selection on community composition, is an appropriate metric in our dataset to measure phylogenetic turnover (Stegen *et al.* 2013).

Within the 21945 community comparisons possible with our data, we found selection to be responsible for 12.3% of the observed community turnover (βNTI>|2|). Most selection increases community similarity (homogeneous selection βNTI<−2: 9.6%), while only 2.7% of community turnover was consistent with heterogeneous selection (βNTI>2). Of the remaining 87.7% variation, 16% was governed by dispersal limitation (βNTI<|2| and RC_Bray_>0.95), whereas 39.1% was attributable to homogenizing dispersal (βNTI<|2| and RC_Bray_<0.95). This leaves 32.6% of community turnover attributable to ecological drift (e.g. stochastic processes) alone (Supp. Fig. 2C). Environmental PCs associated with either selection or drift processes are described in the supplemental material.

## Discussion

Our dataset, encompassing over 200 lakes located across Eastern Canada, showed that watershed anthropogenic activity strongly influences surface water bacterial community composition. Bacterial communities were altered on all levels investigated, ranging from richness, to community composition, to community interactions, to indicator species of high and low lake quality.

### Human impact alters bacterial richness and community composition

The human impact index (HII) variable was found to be associated with surface water communities with significantly lower richness. Specifically, high HII values were associated with high chloride, calcium, magnesium and potassium values, which in turn were associated with reduced richness. Salinity has been shown to be a major factor structuring bacterial diversity in a range of environments (Benlloch *et al.* 2002; Abed *et al.* 2007; Lozupone & Knight 2007), but the effect of salinity on diversity in lake systems is less clear (Wu *et al.* 2006; Wang *et al.* 2011). Importantly, these findings point to a significant impact of salt contamination on the bacterial community even at relatively low levels of salinity (and well below the chronic pollution thresholds for chloride ions at 230 mg/L) and thus suggests that bacterial ecosystem services in lakes may be highly vulnerable to salt contamination.

PC1 was found to be impacted by agriculture, pasture and urban development within the watershed, indicating road salt (including rock salt (NaCl) and other de-icers such as potassium chloride, calcium chloride and magnesium chloride) as a likely source for the environmental variables loaded onto the PC. Chloride concentrations in streams and lakes have been previously shown to be driven by the application of road salt (Kelly *et al.* 2008; Corsi *et al.* 2010) and even relatively low road cover within a watershed (~1%) has been linked to increased chloride concentrations within lakes (Dugan *et al.* 2017).

Community composition was also structured by landscape-scale variables. Lake communities were impacted by nearly every factor explored here, ranging from lake morphometry to most land use classes and spatial components. However, although most factors that we included in our models were significant, our models were only able to explain a small proportion of lake community composition overall (~9-18%). Thus, despite the inclusion of over 30 partially nested variables in the models, we acknowledge that we still fall short of describing the majority of the prokaryotic community diversity observed in this system.

### Human impact weakens bacterial interaction networks

Overall, anthropogenic activity is associated with strong shifts in bacterial community composition. To relate these shifts to community stability and functionality, we investigated how communities in lakes with high, moderate or low human impacted watersheds were altered in their interactions and co-occurrences. As we were interested in overall community structure rather than specific group interactions (e.g. Eiler *et al.* 2012), we treated all interactions (positive and negative) as undirected, to facilitate analysis. Significant co-occurrence may indicate that taxa are directly interacting but may also show taxa as co-occurring due to overall niche similarity (or competitive exclusion in the case of negative co-occurrence). While lake networks constructed from communities from low and moderately impacted watersheds were similar, we detected altered structure and topology in networks from highly impacted lakes. Even though communities from highly impacted lakes had the highest number of taxa involved in significant co-occurrences (220 taxa), the number of significant connections did not scale up proportionally, resulting in, on average, less neighbours per node in these communities than in moderate or low impacted communities. In turn, this resulted in less highly connected keystone taxa and thus a more fragmented network with a lower clustering coefficient (the average number of actual three-node connections passing through each node versus the potential number of three-node connections) and lower network centralization. Higher levels of centralization (i.e. average number of nodes connected to each node or how ‘star-shaped’ a network structure is) have been linked to increased system stability in root microbiomes (Banerjee *et al.* 2019) and lake systems (Peura *et al.* 2015).

Interestingly, moderate HII communities seemed to be only marginally impacted with respect to their co-occurrence network in comparison with low impact communities. This finding may indicate a buffering effect in which microbiomes are able to maintain most ecosystem functions despite shifts in the underlying taxa performing them, for example via functional redundancy. The altered pattern in high impacted communities may indicate that once a threshold is reached, the system’s buffering capacities are exhausted, and that specifically the loss of highly connected and influential keystone taxa may lead to cascading effects of community fragmentation (Peura *et al.* 2015). Alternatively, the patterns could indicate that while low and moderately impacted lakes are quite similar to each other, highly impacted lakes may be overall more variable or that, depending on the exact nature of the anthropogenic impact, multiple high impact communities may exist.

### Ecological and evolutionary processes structuring lake communities

In addition to investigating the effect of watershed anthropogenic activity, the large spatial scale of the study allowed us to assess the ecological and evolutionary processes structuring lake bacterial communities in Eastern Canada in general, as has been previously done in a range of bacterial communities including aquifers (Stegen *et al.* 2013) and the ocean (Logares *et al.* 2018). Firstly, we investigated whether we could identify generalist and specialist taxa within our dataset (Barberán *et al.* 2012). We were able to identify 25 generalist taxa with an occupancy of over 75% in our dataset, including well-known cosmopolitan freshwater lineages such as the actinobacterial *acI* clade ASVs, betaproteobacterial *betI* ASVs and an alphaproteobacterial LD12 ASV. In contrast, we were unable to identify any specialist (low occupancy but relatively high abundance) ASVs, indicating that taxa have either low abundance or wider distributions.

We also utilized the dataset to investigate patterns in community assembly. In accordance with the absence of specialists, we found heterogenizing selection to be only responsible for a very small amount of community variation. Rather, lake ecosystems seemed to be extensively connected by homogenizing dispersal and successful lineages were thus able to establish themselves in other communities (homogenizing selection). Consequently, dispersal limitation was not found to be an important factor structuring lake bacterial communities and instead, ecological drift processes were prevalent. We are not aware of any comparable study of community assembly in lakes, but it was noticeable that, in our system, selection was a much less powerful force shaping communities than in subsurface or marine ecosystems (Stegen *et al.* 2013; Logares *et al.* 2018).

### Caveats and conclusion

Data obtained within the first sampling season of the LakePulse project allows an unprecedented insight into how watershed anthropogenic impact shape bacterial communities on landscape scales. However, the large-scale nature of the project and the sampling effort involved also lead to several caveats. Due to technical difficulties we were unable to measure pH and total phosphorus directly for a large quantity of the lakes and were thus not able to include these variables in our models. pH has been previously shown to be a key driver of bacterial community structure in a range of environments (Lindström *et al.* 2005; Lauber *et al.* 2009; Xiong *et al.* 2012). pH is a complex variable and correlated to some degree with both the concentrations of ions such as chloride and ammonium within the water, as well as lake geography and morphometry. Likewise, phosphate concentrations have been shown to impact lake bacterial community compositions (Tong *et al.* 2005; Romina Schiaffino *et al.* 2011) and sediments (Zeng *et al.* 2009) but were not taken into account in this model. As phosphate, in combination with nitrate, is often used in agricultural settings, we expect some positive correlation between phosphate and nitrogen-species but are unable to directly investigate its role in bacterial community diversity. We suspect that at least some of the unexplained variation in our system is connected to these missing environmental variables. Likewise, we limited our analysis to only include abiotic factors as explanatory variables of bacterial community structure. However, biotic interactions are not only likely to be altered by environmental factors, as shown here via network analysis, but can also exert strong biological control mechanisms for example via anticompetitor compounds (Kraemer *et al.* 2017), predator-prey or mutualistic relationships across trophic levels. Future work within the project is aimed at incorporating bacterial diversity into a food web framework utilizing phytoplankton and zooplankton data also collected from the lakes, as well as deepening our functional understanding of lake communities using metagenomics. Moreover, current sampling efforts are underway that will allow us to include more lakes to extend our frameworks to all of Canada, including regions of extremely high human impact and associated environmental decay due to agriculture and mining.

In conclusion, we used an extensive dataset of lake bacterial communities to model the impact of anthropogenic activity across vast areas of Eastern Canada. Human impact, and specifically variables related to urban and agricultural development within watersheds, had a pronounced effect on lake communities and showed that when high-intensity human activities alter more than about 50% of a watershed, fragmentation of bacterial communities is observed which may ultimately lead to the decline of the ecosystem services provided by them.

## Data accessibility statement

We confirm that the data supporting the results will be made publicly available on Figshare and the data DOI included in the manuscript upon acceptance.

## Acknowledgements

We thank the coordinators and field team members of the LakePulse 2017 sampling campaign for their efforts. We also would like to thank members of the network, and specifically
B. Beisner, V. Fugere and M. Fredette for helpful discussions during the manuscript preparation.

## References

Abed, R.M.M., Kohls, K. & De Beer, D. (2007). Effect of salinity changes on the bacterial diversity, photosynthesis and oxygen consumption of cyanobacterial mats from an intertidal flat of the Arabian Gulf. Environ. Microbiol.

Adrian, R., O’Reilly, C.M., Zagarese, H., Baines, S.B., Hessen, D.O., Keller, W., et al. (2009). Lakes as sentinels of climate change. Limnol. Oceanogr.

Albanese, D., Filosi, M., Visintainer, R., Riccadonna, S., Jurman, G. & Furlanello, C. (2013). Minerva and minepy: A C engine for the MINE suite and its R, Python and MATLAB wrappers. Bioinformatics.

Arbuckle, K.E. & Downing, J.A. (2001). The influence of watershed land use on lake N : P in a predominantly agricultural landscape. Limnol. Oceanogr.

Banerjee, S., Walder, F., Büchi, L., Meyer, M., Held, A.Y., Gattinger, A., et al. (2019). Agricultural intensification reduces microbial network complexity and the abundance of keystone taxa in roots. ISME J.

Barberán, A., Bates, S.T., Casamayor, E.O. & Fierer, N. (2012). Using network analysis to explore co-occurrence patterns in soil microbial communities. ISME J.

Bastviken, D., Cole, J., Pace, M. & Tranvik, L. (2004). Methane emissions from lakes: Dependence of lake characteristics, two regional assessments, and a global estimate. Global Biogeochem. Cycles.

Bates, D., Mächler, M., Bolker, B.M. & Walker, S.C. (2015). Fitting linear mixed-effects models using lme4. J. Stat. Softw.

Benlloch, S., López-López, A., Casamayor, E.O., Øvreås, L., Goddard, V., Daae, F.L., et al. (2002). Prokaryotic genetic diversity throughout the salinity gradient of a coastal solar saltern. Environ. Microbiol.

Callahan, B.J., McMurdie, P.J., Rosen, M.J., Han, A.W., Johnson, A.J.A. & Holmes, S.P. (2016). DADA2: High-resolution sample inference from Illumina amplicon data. Nat. Methods.

Corsi, S.R., Graczyk, D.J., Geis, S.W., Booth, N.L. & Richards, K.D. (2010). A fresh look at road salt: Aquatic toxicity and water-quality impacts on local, regional, and national scales. Environ. Sci. Technol.

Dray, S., Blanchet, G., Borcard, D., Guenard, G., Jombart, T., Legendre, P., et al. (2016). Package “adespatial”: Multivariate Multiscale Spatial Aanalysis.

Dugan, H.A., Bartlett, S.L., Burke, S.M., Doubek, J.P., Krivak-Tetley, F.E., Skaff, N.K., et al. (2017). Salting our freshwater lakes. Proc. Natl. Acad. Sci. U. S. A.

Ecological Stratification Working Group. (1996). A National Ecological Framework for Canada.

Eiler, A., Heinrich, F. & Bertilsson, S. (2012). Coherent dynamics and association networks among lake bacterioplankton taxa. ISME J.

Environment Canada. (2011). 2011 Municipal Water Use Report.

Hair, J., Tatham, R., Anderson, R. & Black, W. (1998). Multivariate Data Analysis. Fifth Edit. Prentice-Hall, London.

Heisler, J., Glibert, P.M., Burkholder, J.M., Anderson, D.M., Cochlan, W., Dennison, W.C., et al. (2008). Eutrophication and harmful algal blooms: A scientific consensus. Harmful Algae.

Hering, D., Borja, A., Carvalho, L. & Feld, C.K. (2013). Assessment and Recovery of European Water Bodies: key Messages from the WISER Project. Hydrobiologia.

Hobbie, S.E., Finlay, J.C., Janke, B.D., Nidzgorski, D.A., Millet, D.B. & Baker, L.A. (2017). Contrasting nitrogen and phosphorus budgets in urban watersheds and implications for managing urban water pollution. Proc. Natl. Acad. Sci.

Huot, Y., Brown, C.A., Potvin, G., Antoniades, D., Baulch, H.M., Beisner, B.E., et al. (2019). The NSERC Canadian Lake Pulse Network: A national assessment of lake health providing science for water management in a changing climate. Sci. Total Environ.

Kara, E.L., Hanson, P.C., Hu, Y.H., Winslow, L. & McMahon, K.D. (2013). A decade of seasonal dynamics and co-occurrences within freshwater bacterioplankton communities from eutrophic Lake Mendota, WI, USA. ISME J.

Kelly, V.R., Lovett, G.M., Weathers, K.C., Findlay, S.E.G., Strayer, D.L., Burns, D.J., et al. (2008). Long-term sodium chloride retention in a rural watershed: Legacy effects of road salt on streamwater concentration. Environ. Sci. Technol.

Kraemer, S.A., Soucy, J.P.R. & Kassen, R. (2017). Antagonistic interactions of soil pseudomonads are structured in time. FEMS Microbiol. Ecol.

Kurtz, Z.D., Müller, C.L., Miraldi, E.R., Littman, D.R., Blaser, M.J. & Bonneau, R.A. (2015). Sparse and Compositionally Robust Inference of Microbial Ecological Networks. PLoS Comput. Biol.

Lauber, C.L., Hamady, M., Knight, R. & Fierer, N. (2009). Pyrosequencing-based assessment of soil pH as a predictor of soil bacterial community structure at the continental scale. Appl. Environ. Microbiol.

Lindström, E.S., Kamst-Van Agterveld, M.P. & Zwart, G. (2005). Distribution of typical freshwater bacterial groups is associated with pH, temperature, and lake water retention time. Appl. Environ. Microbiol.

Logares, R., Tesson, S.V.M., Canback, B., Pontarp, M., Hedlund, K. & Rengefors, K. (2018). Contrasting Prevalence of Selection and Drift in the Community Structuring of Bacteria and Microbial Eukaryotes. Environ. Microbiol. 20, 2231–2240.

Lozupone, C.A. & Knight, R. (2007). Global patterns in bacterial diversity. Proc. Natl. Acad. Sci., 104, 11436–11440.

Marmen, S., Blank, L. Al-Ashhab, A., Malik, A., Ganzert, L. Lalzar, M., Grossart, H.-P. & Sher, D. (2019). The Role of Land Use Types and Water Chemical Properties in Structuring the Microbiome of a Connected lake System. bioRxiv.

Martin, M. (2011). Cutadapt removes adapter sequences from high-throughput sequencing reads. EMBnet.journal.

McMurdie, P.J. & Holmes, S. (2013). Phyloseq: An R Package for Reproducible Interactive Analysis and Graphics of Microbiome Census Data. PLoS One.

Novotny, E. V., Murphy, D. & Stefan, H.G. (2008). Increase of urban lake salinity by road deicing salt. Sci. Total Environ.

Oksanen, J., Blanchet, F.G., Friendly, M., Kindt, R., Legendre, P., Mcglinn, D., et al. (2016). Vegan: Community Ecology Package. URL https://cran.r-project.org, https://github.com/vegandevs/vegan.

Patton, C. & Kryskalla, J. (2003). Methods of Analysis by the U.S. Geological Survey National Water Quality Laboratory - Evaluation of Alakline Digestion as an Alternative to Kjedahl Digestion for Determination of Total and Dissolved Nitrogen and Phosphorous. Water-Resources Investig., 03.

Peura, S., Bertilsson, S., Jones, R.I. & Eiler, A. (2015). Resistant microbial cooccurrence patterns inferred by network topology. Appl. Environ. Microbiol.

Pinheiro, J., Bates, D., DebRoy, S., Sarkar, D. & R Development Core Team, T. (2019). nlme: Linear and Nonlinear Mixed Effect Models. R package version 3.1–141.

Price, M.N., Dehal, P.S. & Arkin, A.P. (2010). FastTree 2 - Approximately maximum-likelihood trees for large alignments. PLoS One.

Quast, C., Pruesse, E., Yilmaz, P., Gerken, J., Schweer, T., Yarza, P., et al. (2013). The SILVA ribosomal RNA gene database project: Improved data processing and web-based tools. Nucleic Acids Res.

Rohwer, R.R., Hamilton, J.J., Newton, R.J. & McMahon, K.D. (2018). TaxAss: Leveraging a Custom Freshwater Database Achieves Fine-Scale Taxonomic Resolution. mSphere.

Romina Schiaffino, M., Unrein, F., Gasol, J.M., Massana, R., Balagué, V. & Izaguirre, I. (2011). Bacterial community structure in a latitudinal gradient of lakes: The roles of spatial versus environmental factors. Freshw. Biol.

Rosseel, Y. (2012). Lavaan: An R package for structural equation modeling. J. Stat. Softw.

Scavia, D., David Allan, J., Arend, K.K., Bartell, S., Beletsky, D., Bosch, N.S., et al. (2014). Assessing and addressing the re-eutrophication of Lake Erie: Central basin hypoxia. J. Great Lakes Res.

Shade, A., Kent, A.D., Jones, S.E., Newton, R.J., Triplett, E.W. & McMahon, K.D. (2007). Interannual dynamics and phenology of bacterial communities in a eutrophic lake. Limnol. Oceanogr., 52, 487–494.

Stegen, J.C., Lin, X., Fredrickson, J.K., Chen, X., Kennedy, D.W., Murray, C.J., et al. (2013). Quantifying community assembly processes and identifying features that impose them. ISME J.

Taranu, Z.E. & Gregory-Eaves, I. (2008). Quantifying relationships among phosphorus, agriculture, and lake depth at an inter-regional scale. Ecosystems.

Tong, Y., Lin, G., Ke, X., Liu, F., Zhu, G., Gao, G., et al. (2005). Comparison of microbial community between two shallow freshwater lakes in middle Yangtze basin, East China. Chemosphere.

Tranvik, L.J., Downing, J.A., Cotner, J.B., Loiselle, S.A., Striegl, R.G., Ballatore, T.J., et al. (2009). Lakes and reservoirs as regulators of carbon cycling and climate. Limnol. Oceanogr., 54, 2298–2314.

U.S. Environmental Protection Agency. (1994). Method 200.7: Determiantion of Metals and Trace Eelements in Water and Wastes by Inductively Coupled Plasma-Atomic Emission Spectrometry.

U.S. Environmental Protection Agency. (1997). Method 300.1: Determination of Inorganic Anions in Drinking Water by Ion Chromatography.

U.S. Environmental Protection Agency. (2009). National Lake Assessment: A Collaborative Survey of the Nation’s Lakes. EPA 841-R-09-001.

Wang, J., Yang, D., Zhang, Y., Shen, J., van der Gast, C., Hahn, M.W., et al. (2011). Do Patterns of Bacterial Diversity along Salinity Gradients Differ from Those Observed for Macroorganisms? PLoS One.

Wu, Q.L., Zwart, G., Schauer, M., Kamst-Van Agterveld, M.P. & Hahn, M.W. (2006). Bacterioplankton community composition along a salinity gradient of sixteen high-mountain lakes located on the Tibetan Plateau, China. Appl. Environ. Microbiol.

Wu, Y. (2014). Barcode Demultiplex for Illumina I1, R1, R2 fastq.gz files.

Xiong, J., Liu, Y., Lin, X., Zhang, H., Zeng, J., Hou, J., et al. (2012). Geographic distance and pH drive bacterial distribution in alkaline lake sediments across Tibetan Plateau. Environ. Microbiol.

Zeng, J., Yang, L., Li, J., Liang, Y., Xiao, L., Jiang, L., et al. (2009). Vertical distribution of bacterial community structure in the sediments of two eutrophic lakes revealed by denaturing gradient gel electrophoresis (DGGE) and multivariate analysis techniques. World J. Microbiol. Biotechnol.

